# Calcium-Activated Sarcomere Contractility Drives Cardiomyocyte Maturation and the Response to External Mechanical Cues but is Dispensable for Sarcomere Formation

**DOI:** 10.1101/2025.03.18.644054

**Authors:** Laura A. Sherer, Abigail Nagle, Mary Papadaki, Seby Edassery, Dasom Yoo, Lauren D’Amico, Daniel Brambila-Diaz, Michael Regnier, Jonathan A. Kirk

**Author notes:** Contributed Equally to This Work (Laura A. Sherer, Abigail Nagle). Corresponding Author Jonathan A. Kirk, Ph.D. Department of Cell and Molecular Physiology Loyola University Chicago Stritch School of Medicine Center for Translational Research, Room 522 2160 S. First Ave. Maywood, IL 60153.

## Abstract

**Background:** Understanding the mechanisms of cardiomyocyte development is critical for fulfilling the potential of induced pluripotent stem cell-derived cardiomyocytes (iPSC-CMs). Although myocyte development is known to depend on internal and external mechanical cues, further investigation is required to understand the contributions of different signals and how they are integrated together to generate an adult cardiomyocyte. Here, we address this gap by examining the role of calcium-activated contractility in sarcomere formation and maturation and its influence on the iPSC-CM response to nanopatterns.

**Methods:** We generated iPSCs with homozygous D65A cardiac troponin C (cTnC) mutations. This mutation prevents calcium binding to site II of cTnC, resulting in tropomyosin blocking strong myosin binding to the thin filament and inhibiting sarcomere contraction. The iPSCs were differentiated into cardiomyocytes and matured in culture over 60 days. Cells were characterized via fluorescence imaging and calcium transient analysis. WT and mutant proteomes were examined via mass spectrometry throughout differentiation and maturation. We also replated partially matured cardiomyocytes onto nanopatterned surfaces to investigate how external mechanical signals affect maturation in contractile versus non-contractile cells.

**Results:** Surprisingly, we found that sarcomeres formed in the cTnC D65A cardiomyocytes, though these sarcomeres were underdeveloped and disorganized. Mutant cardiomyocytes also exhibited significant proteomic maturation defects and abnormal calcium transients. Plating D65A cardiomyocytes on nanopatterns improved structural and proteomic maturation. However, plating WT cardiomyocytes on nanopatterns led to a reduction in sarcomeric and oxidative phosphorylation protein content.

**Conclusions:** Calcium-activated contractility is dispensable for sarcomerogenesis but critical for cardiomyocyte maturation. In non-contractile, mutant cardiomyocytes, nanopatterns enhance maturation, suggesting that external mechanical cues may partially compensate for defective contractility. However, nanopatterns did not facilitate WT maturation and may have hindered it. In addition to these novel findings, these large mass spectrometry datasets cataloging iPSC-CM maturation represent a useful resource for the cardiovascular community.

## Introduction

Cardiomyocyte development is characterized by extensive structural, metabolic, and electrophysiological changes. Cardiac sarcomeres form early in mammalian embryos, with contractile activity detected in humans at Day 16 post-fertilization.^1^ Sarcomere assembly is followed by cardiomyocyte maturation. During maturation, cardiomyocytes exit the cell cycle, grow larger, and elongate.^2,3^ They develop and/or expand cellular infrastructure like t-tubules and the sarcoplasmic reticulum to facilitate calcium handling. Their myofibrils grow and become more uniaxially organized, allowing them to contract with progressively more force. To meet their increasing metabolic demands, maturing cardiomyocytes upregulate oxidative phosphorylation (OXPHOS), specifically fatty acid oxidation. The proper execution of this maturation program depends on the cardiomyocyte’s ability to respond to and coordinate both external and internal mechanical signals. Failure of cardiomyocytes to enact these changes is associated with congenital heart diseases and heart failure,^4^ and, therefore, determining the role and relative contributions of different mechanical cues in cardiomyocyte development could facilitate therapeutic development.

Understanding the mechanisms of cardiomyocyte development has become increasingly important with the emergence of human induced pluripotent stem cell (iPSC)-derived cardiomyocytes (iPSC-CMs). iPSCs can be patient specific, making them incredibly useful for applications like drug screening, disease modeling, and ultimately, patient tissue grafts.^2^ However, a drawback of iPSC-CMs is that they are more similar to perinatal cardiomyocytes rather than adult cardiomyocytes.^5^ iPSC-CMs are smaller and rounder with less developed, weaker sarcomeres and slower calcium dynamics than their adult counterparts.^2,3^

There are several ways to ameliorate aspects of iPSC-CM maturation. These interventions include co-culture with other cell types, altered media composition, nanopatterned surfaces, engineered heart tissues, electrical stimulation, and prolonged culture.^2,3^ While each of these methods enhance aspects of maturation, none produce fully mature myocytes, underscoring the complex nature of maturation and the numerous external and internal signals that must be coordinated to generate an adult cardiomyocyte. An understanding of the interplay between these different cues is crucial for driving iPSC-CM maturation forward.

An advantage of the immaturity of iPSC-CMs is that they can disassemble and reassemble their sarcomeres. Researchers have leveraged this characteristic to gain insight into sarcomere assembly.^6,7^ Recent evidence supports the pre-myofibril model, which has become the prevailing idea of sarcomere formation.^6–9^ In this model, muscle-specific stress fibers form at the edges of cardiomyocytes. The stress fibers transition into myofibrils as they migrate into the cell body, potentially acting like a template for sarcomere formation.

Despite recent advances, many aspects of sarcomerogenesis are unresolved. For example, the role of calcium-activated contractility in sarcomere assembly is still unclear, though loss of contractility is associated with maturation defects.^10^ Inhibition of myo-II or subsets of myo-II in different model systems appear to both prohibit and permit sarcomere formation.^7,10^ These studies are difficult to interpret because acute pharmacological inhibition can result in incomplete inhibition or off-target effects. As such, the role of contractility in sarcomere formation needs to be clarified.

In this study, we investigated how iPSC-CMs respond to and integrate internal and external mechanical cues during cardiomyocyte development. Specifically, we examined the role of contractility (i.e., an internal cue) on sarcomere formation and maturation using a gene editing approach to introduce the D65A mutation into cardiac troponin C (cTnC). This eliminates calcium-mediated signaling for thin filament activation, blocking troponin-dependent myosin binding to actin to form crossbridges and crossbridge cycling activity. Our results show that although the D65A mutation caused significant maturation defects, cardiomyocytes still formed sarcomeres, establishing the dispensability of contractility in sarcomere assembly. We also assessed the interaction between nanopatterns (i.e., an external cue) and sarcomere contractility (an internal cue) during cardiomyocyte maturation. Nanopatterns enhanced structural and proteomic maturation metrics in the non-contractile D65A cardiomyocytes. However, for WT cardiomyocytes, nanopatterns decreased sarcomeric and OXPHOS protein expression, potentially suggesting disrupted WT maturation. In addition, we have generated multiple large mass spectrometry datasets that catalog iPSC-CM proteomic maturation and have made protein abundance time courses easily accessible, thus providing a useful resource for the cardiovascular community.

## Methods

Detailed methods are available in the **Supplemental Materials**. Proteomics data is available at ftp://massive.ucsd.edu/v07/MSV000096590/. (Reviewers, please use ftp://MSV000096590@massive.ucsd.edu, password: g01P4kp822bMXr16).

### CRISPR/Cas9 Targeting of TNNC1 in hiPSCs

Homozygous TNNC1 D65A (denoted cTnC D65A) was generated using CRISPR-Cas9 by the Tom & Sue Ellison Stem Cell Core at the Institute for Stem Cell & Regenerative Medicine at the University of Washington and confirmed via sequencing (**Figure S1A-B**).

### iPSC-Cardiomyocyte Generation and Maturation Protocol

iPSCs were differentiated into cardiomyocytes in a biphasic Wnt modulation protocol adapted as previously described.^11^ Cells were purified in glucose-free DMEM with 4 mM sodium L-lactate from Days 16-20. From Day 20 onward, cells were fed with cardiomyocyte maintenance media (RPMI containing 1X B27 supplement with insulin), with media changes every 2-3 days. iPSC-CM imaging and calcium transient analysis were carried out as described in the Supplemental Methods.^12–14^

### Mass Spectrometry and Analysis

Wildtype and cTnC D65A cells were flash frozen on Days 5, 7, 14, 30, and 60 for mass spectrometry analysis (n = 3 differentiations per group). Samples were digested with Trypsin/LysC, desalted using Sep-Pak tC18 cartridges, and analyzed via LC/MS-MS using a c18 PrepMap column and an Orbitrap Eclipse Tribrid mass spectrometer. The raw MS data was analyzed using Proteome Discoverer (2.5, Thermo Fisher). NIH’s BRB-ArrayTools software was also used for a time course analysis. Proteins that were significantly different over time were compiled into a spreadsheet with links to graphs of protein abundances as a function of time. This data is available at https://github.com/JKirkLab/iPSCProteomics. Gene ontology (GO) term analysis was carried out using DAVID bioinformatics resources.^15,16^ Graphs were plotted in Prism 10.

### Statistical Analysis

Proteomics statistics and GO term p-values were obtained using Proteome Discoverer and the DAVID web server, respectively. For imaging and calcium transient analysis, Prism 10 was used to perform unpaired t-tests or a 2-way ANOVA with multiple comparisons.

## Results

### cTnC D65A Cardiomyocytes Form Disorganized and Underdeveloped Myofibrils

To investigate the role of internal mechanical cues on the maturation of iPSC-derived cardiomyocytes (iPSC-CMs), CRISPR-Cas9 was used to generate iPSCs with homozygous cardiac Troponin C (cTnC) D65A mutations (confirmed by sequencing, **Figure S1)**. This mutation prevents cTnC from binding to calcium in the N-terminal regulatory site II.^17^ As a result, the troponin complex does not undergo the conformational change that allows tropomyosin to move away from myosin binding sites on the thin filament, preventing sarcomere contraction.

Cardiomyocytes were differentiated and matured for up to 60 days (**Figure 1A**). Wildtype (WT) iPSC-CMs first exhibited spontaneous contractions at approximately Day 7, whereas beating was never observed in the D65A cardiomyocytes. Cells were replated at Day 14 and metabolically purified in glucose-free media containing lactate from Days 16 to 20. Purified cardiomyocytes were cultured in standard RPMI media supplemented with B27 until Day 60. Cells were characterized via imaging, calcium transient analysis, and mass spectrometry at different timepoints throughout this process.

**Figure 1.**
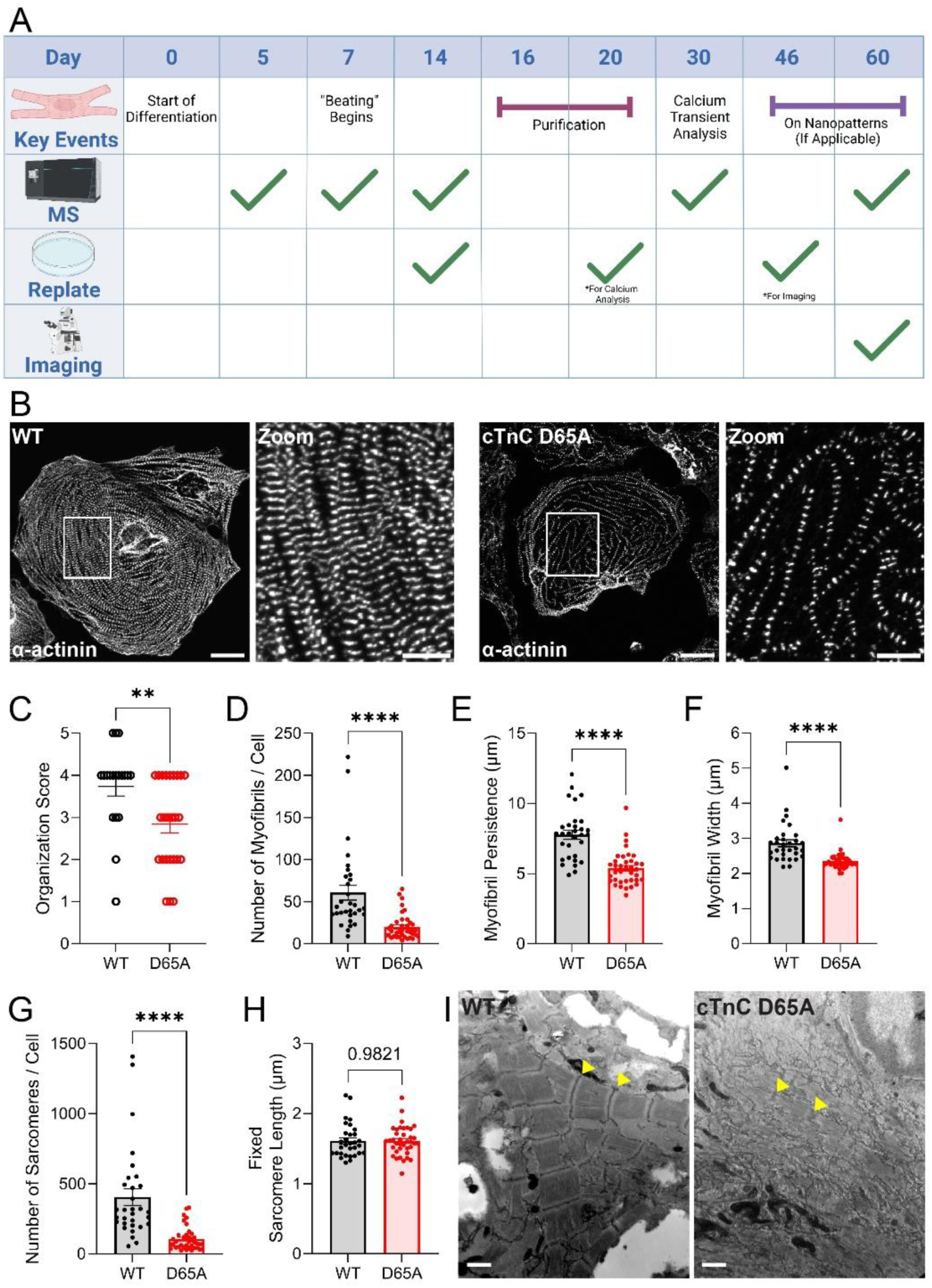
cTnC D65A cardiomyocytes develop disorganized sarcomeres. **A)** Schematic outlining the experimental timeline. Created in BioRender. **B)** Human iPSC-derived cardiomyocytes were labeled with α-actinin-2 antibody and imaged with fluorescence microscopy. Representative images are shown. Scale bars = 30 µm for whole cell images, 10 µm for zoomed images. **C)** Organization scores for wildtype (WT) and cTnC D65A cardiomyocytes quantified from α-actinin-labeled micrographs. **D-H)** Automated quantification of the number of myofibrils per cell **(D)**, myofibril persistence (the length over which myofibrils remain straight) **(E)**, myofibril width **(F)**, number of sarcomeres per cell **(G)**, and sarcomere length **(H)** as determined from cells fixed in blebbistatin and labeled with α-actinin antibody. Asterisks (*<0.05, **<0.01, ***<0.001, ****<0.0001) indicate significance as determined by Welch’s t-test. **I)** Transmission electron micrographs of WT and cTnC D65A cardiomyocytes. Yellow arrows point to adjacent z-discs. Scale bar = 1 µm.

To evaluate sarcomere structure on Day 60, we labeled the cardiomyocytes with α-actinin-2 antibody, which localizes strongly to z-discs, and imaged via fluorescence microscopy (**Figure 1B**). The striated pattern of sarcomeres was clearly visible in both WT and D65A cardiomyocytes. However, mutant sarcomeres appeared less abundant and more disordered, clearly lacking higher-order structure. Sarcomere organization was blindly scored on a scale of 1 to 5, with 1 representing sparse, punctate z-discs and 5 corresponding to highly regular, aligned z-discs covering most of the cell area.^13^ D65A cardiomyocytes had a significantly lower organization score compared to wildtype (**Figure 1C**). The number of myofibrils per cell (**Figure 1D**), myofibril persistence (the length over which myofibrils remain straight) (**Figure 1E**), myofibril width (**Figure 1F**), and number of sarcomeres per cell (**Figure 1G**) were also significantly reduced in the mutant myocytes. Sarcomere length (**Figure 1H**), measured in the presence of blebbistatin to inhibit myosin binding to actin, was unchanged. We also imaged the cardiomyocytes via transmission electron microscopy (**Figure 1I**). D65A sarcomeres were underdeveloped and exhibited wavy z-discs, supporting our immunofluorescence data. Thus, the absence of calcium-activated force generation does not abolish sarcomere formation but does lead to smaller and more disordered myofibrils.

To assess differences in the dynamics of myofibril formation, we replated mCherry-α-actinin-expressing WT and mutant cells onto glass and imaged while they underwent myofibril re-assembly (**Movie S1-S2**). Whereas WT exhibited the canonical centripetal formation of myofibrils,^7^ D65A cardiomyocytes myofibrils failed to add sarcomeres to existing myofibrils.

To investigate how loss of sarcomere contraction via the cTnC D65A mutation changes the proteome over time, whole cell samples were collected at Days 5, 7, 14, 30 and 60 and analyzed using mass spectrometry (**Figure 1A, Table S1**). Time course analysis of protein abundances is available at https://github.com/JKirkLab/iPSCProteomics as a resource for the cardiovascular research community.

To understand broad patterns within the dataset, we performed principal component analysis (PCA) (**Figure 2A, Table S2**). Strikingly, both WT and D65A samples cluster based on time along the first principal component (PC1), with positive PC1 values associated with later timepoints. Given that PC1 represents a sizable portion (31.3%) of the dataset variability, these patterns suggest that many time-dependent proteomic changes are consistent between WT and D65A cardiomyocytes. In contrast, along principal component 2 (PC2,12.7%), WT and D65A exhibit inverted trajectories, with increasing PC2 values over time for D65A and decreasing values for WT. Thus, PC2 may reflect proteome changes that differ between WT and D65A during maturation.

**Figure 2.**
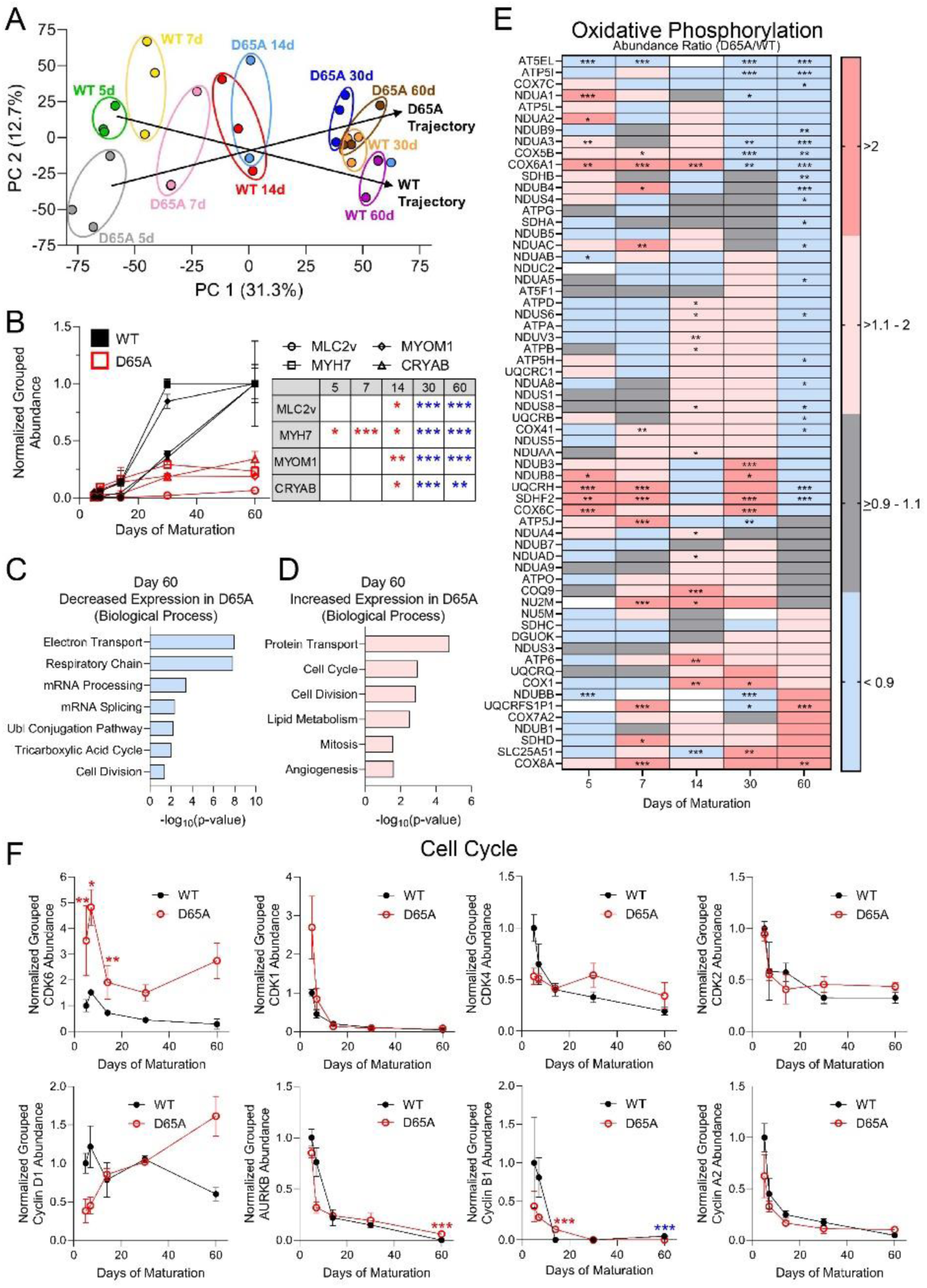
cTnC D65A cardiomyocytes have reduced expression of maturation markers. Protein lysates from WT and cTnC D65A cardiomyocytes were analyzed by mass spectrometry to obtain grouped abundances and abundance ratios (D65A/WT). Asterisks specify p-values (*<0.05, **<0.01, ***<0.001) as determined by Proteome Discoverer. Color specifies higher (red) or lower (blue) abundance in mutant compared to WT. Error bars show the standard error of the mean. **A)** Principal component analysis of WT and D65A mutant at different timepoints. Individual samples are represented by a dot. Colors and ovals indicate samples from the same group. **B)** Grouped abundances for WT (black) and mutant (red) maturation markers over time. Values for each protein were normalized to WT grouped abundance on Day 60. **C-D)** Biological Process Gene Ontology (GO) terms enriched among proteins that were significantly downregulated **(C)** or upregulated **(D)** in the D65A mutant compared to WT on Day 60. **E)** Heat map of abundance ratios (D65A/WT) for oxidative phosphorylation proteins throughout maturation. White denotes proteins that were not detected in the WT or the mutant. **F)** Grouped abundances for cell cycle regulators over time in WT (black) and cTnC D65A (red) cardiomyocytes.

### cTnC D65A Cardiomyocytes Exhibit Maturation Defects

To better understand how D65A influences proteomic maturation, we evaluated the abundance of 4 robust maturation markers^2,3,18,19^ (CRYAB, MYH7, MLC2v, and MYOM1). All 4 markers increased with time in the WT cardiomyocytes (**Figure 2B**), confirming enhanced maturation with longer culture times and validating our maturation protocol. There was also a slight increase of these markers in D65A cardiomyocytes, suggesting some degree of maturation progression. Nevertheless, all 4 markers were significantly lower in the D65A mutant compared to the WT from Day 30 onward. These data imply that the internal mechanical cues associated with sarcomere contraction facilitate cardiomyocyte maturation.

We next sought to identify what specific aspects of maturation are disrupted by cTnC D65A, so we conducted gene ontology (GO) enrichment analysis (**Figure 2C-D**). Proteins with decreased expression in the mutant versus WT cardiomyocytes at Day 60 were enriched in electron transport and respiratory chain proteins (**Figure 2C**). We followed up by analyzing a panel of proteins involved in oxidative phosphorylation (OXPHOS). As expected, OXPHOS proteins generally increased in abundance over time in the WT (**Figure S2**), reflecting the characteristic upregulation in OXPHOS during normal maturation. Conversely, D65A cardiomyocytes typically downregulated OXPHOS proteins between Days 30 and 60.

Out of 62 OXPHOS proteins, many (∼48%) were significantly elevated in the mutant over the WT at one or more of the early timepoints (Days 5-30) (**Figure 2E**). Given that metabolic purification occurs at Days 16-20, it is possible that some of the increase in OXPHOS protein content at Days 5, 7, and 14 comes from the non-cardiomyocyte cell population that arises during the differentiation protocol. However, because cTnC is only expressed in cardiomyocytes and the mutation does not appear to alter differentiation efficiency (**Figure S3**), the difference between wildtype and mutant likely arises from cardiomyocytes or cardiomyocyte progenitors. These early, elevated OXPHOS protein abundances in the D65A cells may be part of an initial compensatory response caused by loss of internal mechanical cues. In contrast, on Day 60, ∼34% of OXPHOS proteins were significantly decreased in the D65A cardiomyocytes compared to WT, whereas only ∼3% were significantly upregulated. Thus, the proteomics of D65A cardiomyocytes suggest a reduced reliance on OXPHOS to meet metabolic needs on Day 60.

Cardiomyocyte maturation is associated with a decline in proliferation. Proteins that were differentially expressed between D65A and WT cardiomyocytes were often enriched with GO-term annotations like “mitosis”, “cell division”, and “cell cycle” (**Figure 2C-D**, **Figure S4A**). However, it is difficult to interpret this finding by itself because these annotations do not distinguish between proteins that arrest or promote cell cycle progression. It also does not consider a protein’s degree of importance in these processes—whether they are critical or marginal. To clarify how cell cycle regulation is altered in the D65A mutant, we examined the abundance of established mitotic regulators like AURKB, CDK1/2/4/6, and cyclins (A2, B1, D1) (**Figure 2F**). These proteins drive proliferation and are repressed during maturation.^3,20–22^ As expected, 7 out of 8 of these cell cycle proteins (CDK6, CDK1, CDK4, CDK2, AURKB, Cyclin B1, Cyclin A2) exhibit a general downward trajectory over time in both the D65A mutant and wildtype, suggesting that both groups become less proliferative as they mature.

Calcium transient amplitudes increase during maturation, and the release and reuptake of calcium ions from/to the sarcoplasmic reticulum (SR) becomes faster and more efficient. We investigated how the D65A mutation alters calcium transients using the IonOptix system (**Figure 3A**). On Day 30, the WT and D65A cardiomyocytes showed no significant change in baseline calcium concentration (**Figure 3B**). However, D65A cardiomyocytes did have increased peak calcium release relative to baseline (**Figure 3C**) and exhibited faster times to peak calcium levels (**Figure 3D**) and slower times to 10% baseline levels (**Figure 3E**).

**Figure 3.**
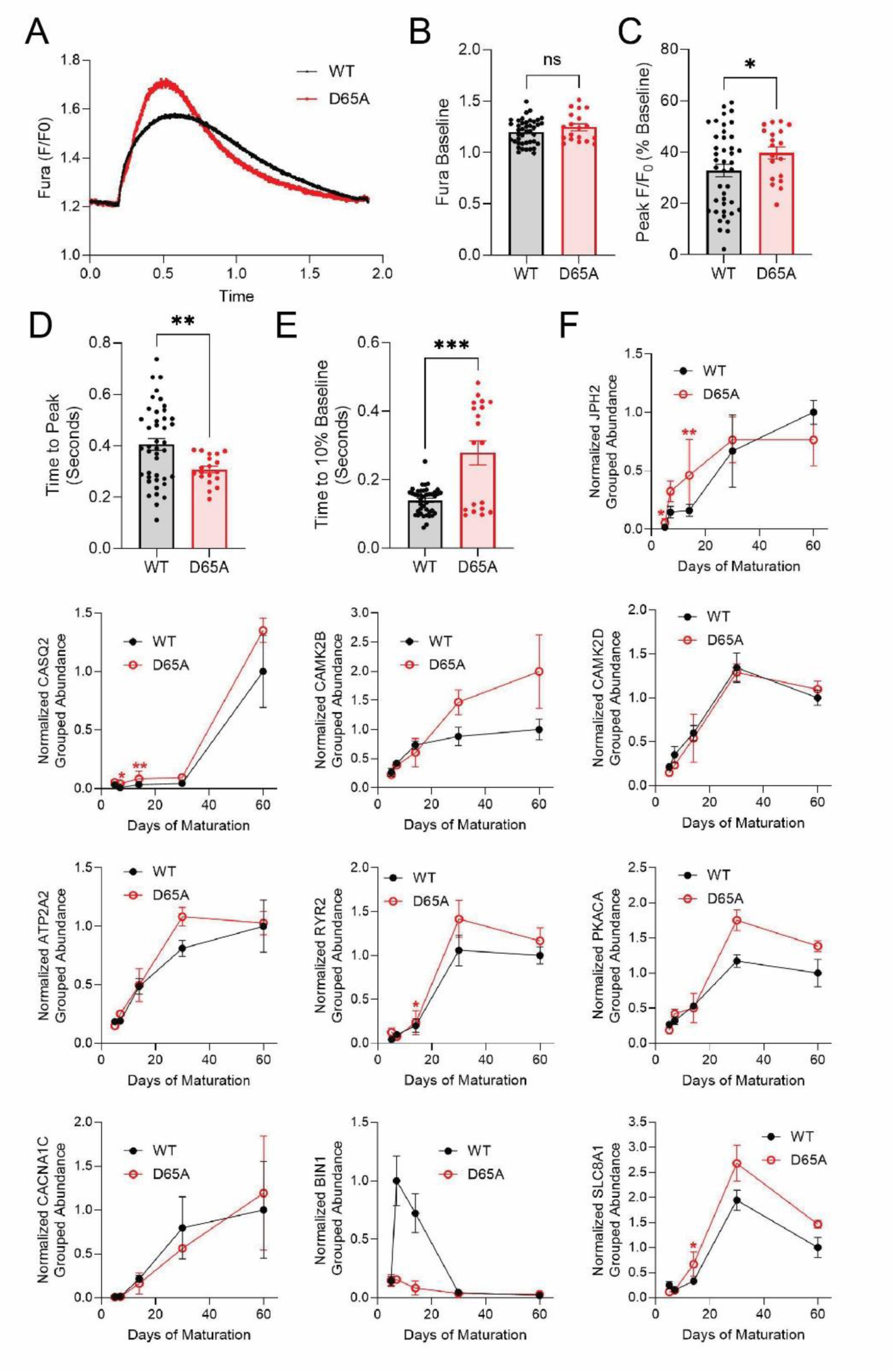
Calcium transients are altered in cTnC D65A cardiomyocytes. **A)** Average fura-2AM IonOptix calcium flux trace of WT and D65A cardiomyocytes. **B-E)** Quantification of calcium flux data from fura-2AM IonOptix for baseline calcium concentration **(B)**, peak calcium release normalized to baseline **(C),** time to peak **(D)**, and time to 10% baseline **(E)**. **F)** Grouped abundances for calcium handling proteins in WT (black) and cTnC D65A (red) cardiomyocytes over time, as determined by mass spectrometry. Asterisks indicate significant upregulation (red) or downregulation (blue) in the mutant with respect to WT. P-values (*<0.05, **<0.01, ***<0.001) were calculated by Proteome Discoverer.

We next determined whether these differences in calcium transients were mediated by changes in calcium handling protein abundance. We observed a general increase in calcium handling proteins during maturation in both the WT and cTnC D65A cardiomyocytes (**Figure 3F**). The only exception to this upward trend was BIN1, a protein associated with T-tubule development. Generally, the differences in calcium handling protein abundance between the WT and mutant were not significant. However, when a statistically significant difference existed, the D65A cardiomyocytes expressed more of the calcium handling protein than the WT. These proteomic differences and altered calcium transients likely result from a large decrease in calcium buffering induced by loss of calcium binding to site II of cTnC, which is the cell’s largest calcium buffer.^23^

Since an increase in sarcomere content is critical to cardiomyocyte maturation, we investigated changes to sarcomere protein abundance over time. Most sarcomere proteins were significantly reduced in the mutant compared to the WT at Days 30 and 60 (**Figure 4A-B, Figure S5**), suggesting underdeveloped myofibrils, in agreement with our structural data (**Figure 1**). In contrast, many sarcomere proteins were significantly elevated at earlier timepoints in the D65A mutant. Like for OXPHOS proteins, these increases could represent a compensatory response to deficient sarcomere formation and/or contraction. We also observed that D65A cells failed to undergo the characteristic isoform switch from MYH6 to MYH7 during maturation.

**Figure 4.**
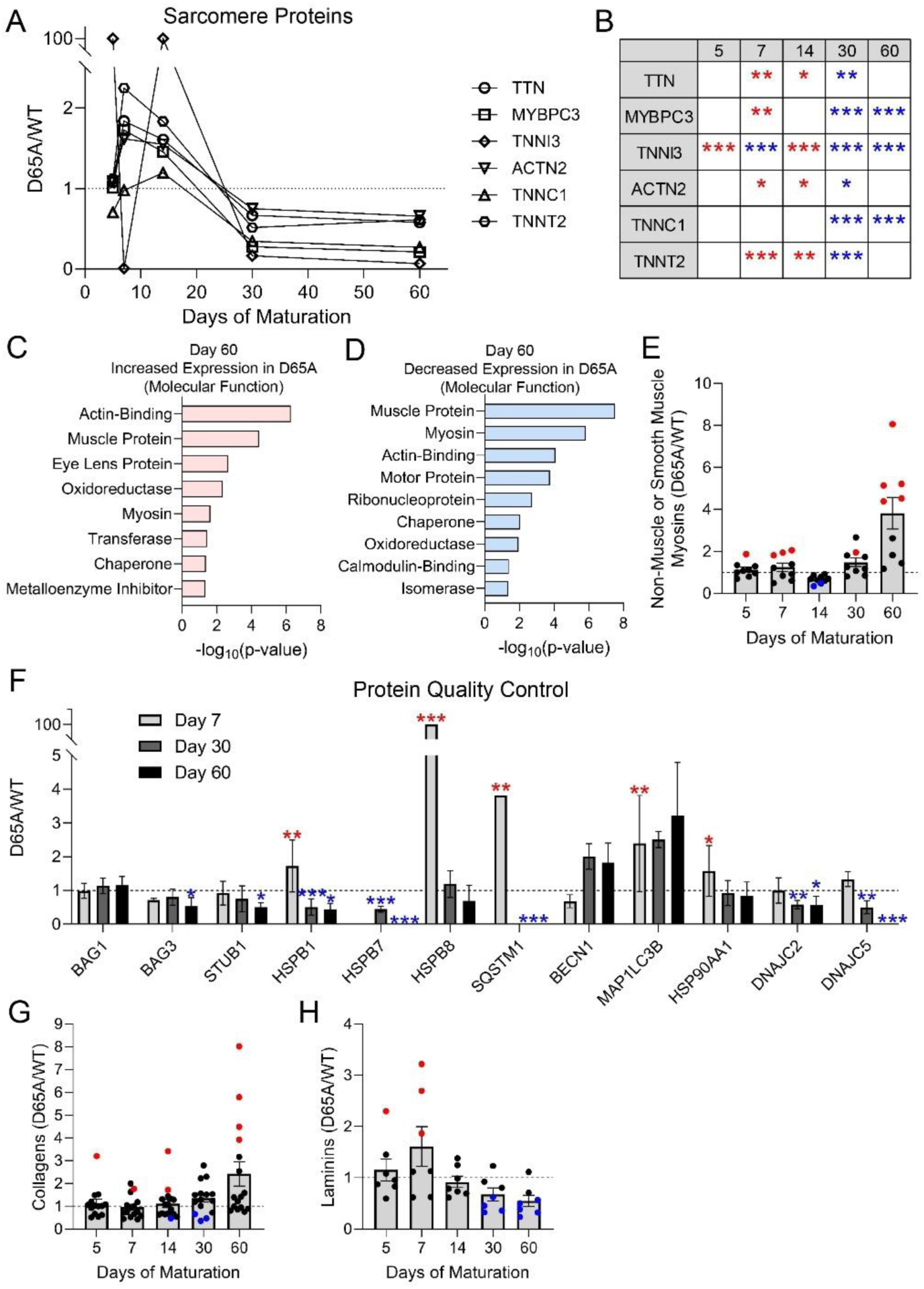
cTnC D65A cardiomyocytes exhibit aberrant expression patterns for sarcomeric proteins, protein quality control machinery, and extracellular matrix proteins. **A-B)** Select sarcomeric protein abundance ratios (D65A/WT) **(A)** and their p-values **(B)** over time for WT and cTnC D65A cardiomyocytes. Dotted line marks an abundance ratio of 1. **C-D)** Molecular Function GO terms enriched among proteins significantly increased **(C)** or decreased **(D)** in cTnC D65A cardiomyocytes compared to WT on Day 60. **E)** Abundance ratios for non-muscle myosins throughout maturation. **F)** Abundance ratios for protein quality control machinery (chaperones, co-chaperones, autophagy proteins) on Days 7, 30, and 60. **G-H)** Abundance ratios (D65A/WT) for panels of collagens **(G)** or laminins **(H)**. Significant differences between WT and mutant are indicated by asterisks (*<0.05, **<0.01, ***<0.001) or colored dots. Color specifies higher (red) or lower (blue) abundance in the mutant compared to WT. Error bars show the standard error of the mean.

We performed Molecular Function GO-term enrichment analyses to identify other changes induced by the D65A mutation (**Figure 4C-D**). As expected, proteins differentially expressed at Day 60 were enriched in actin-binding proteins, muscle proteins, and myosins. Myosins with increased abundance in D65A cardiomyocytes were overwhelmingly non-muscle or smooth muscle myosins (MYH10, MYH11, MYL6, MYL9)^24,25^(**Figure 4E**), whereas myosins depressed in D65A samples (MYH7, MYH1) were linked to cardiac and skeletal muscle.^25^

### cTnC D65A Cardiomyocytes Have Altered Expression of Protein Quality Control Machinery and Extracellular Matrix Proteins

Since the differentially expressed proteins were enriched in chaperones at multiple timepoints (**Figure 4C**, **Figure S4B**), we predicted that protein quality control (PQC) systems were altered in D65A cardiomyocytes. We assembled a panel of PQC proteins that includes components of the CASA (chaperone-assisted selective autophagy) complex, like BAG3, STUB1, and HSPB8. The CASA pathway is upregulated in striated muscle and in response to mechanical stress,^26^ mediating turnover of numerous sarcomere proteins.^27^ Because contraction is a major driver of protein damage, the loss of sarcomere contractility in the D65A cardiomyocytes could potentially reduce the requirement for CASA, making CASA components compelling proteins to examine. Our panel also includes other autophagy proteins (BECN1, MAP1LC3B, SQSTM1), chaperones (HSP90AA1, HSPB1, HSPB7), and co-chaperones (BAG1, DNAJC2, DNAJC5). On Day 7, D65A cardiomyocytes had elevated levels of several PQC proteins (HSPB1, HSPB8, MAP1LC3B, and SQSTM1) (**Figure 4F**), implying increased cellular stress. In contrast, on Day 60, many PQC proteins (BAG3, STUB1, HSPB7, SQSTM1, DNAJC2, and DNAJC5) were less abundant in the D65A cells than the WT. This finding suggests that loss of sarcomere contraction reduces the demand for PQC machinery like the CASA complex.

Myocardial ECM is mostly deposited by cardiac fibroblasts, but cardiomyocytes also produce ECM constituents like collagens and laminins.^28,29^ It has been established that ECM composition influences both cardiomyocyte maturation and contractility^30–33^ and that the ECM composition changes during development and disease.^33,34^ Less is known about how sarcomere contractility and maturation affect ECM deposition by cardiomyocytes. To assess this, we examined the abundance of different collagen and laminin isoforms. Several collagens (COL1A1, COL1A2, COL2A2, COL14A1) were drastically (>300%) more abundant in D65A cardiomyocytes than WT on Day 60 (**Figure 4G**). In contrast, 5 out of the 7 laminins in our panel were significantly reduced (<60%) (**Figure 4H**). This data demonstrates that the absence of contractility during maturation produces distinct changes in cardiomyocyte-mediated ECM deposition and suggests differences in ECM sensing between mutant and WT.

### Nanopatterns Promote Structural Maturation of cTnC D65A Cardiomyocytes

We next sought to understand the interaction between internally generated force (i.e., sarcomere contraction) and external mechanical cues. WT and D65A cardiomyocytes were replated onto either flat or nanopatterned surfaces on Day 46. To determine structural differences between mutant and WT, we stained the cells with phalloidin-488, which binds to actin filaments, and imaged on Day 60 (**Figure 5A**). D65A cardiomyocytes were more elongated on the nanopatterned surface (**Figure 5B**). As previously seen (**Figure 1**), mutant cardiomyocytes on flat surfaces had fewer sarcomeres and myofibrils per cell (**Figure 5C-D**) and shorter, thinner myofibrils (**Figure 5E-F**) compared to WT. Interestingly, these differences between WT and D65A were abolished on nanopatterns. D65A cardiomyocytes also exhibited increased myofibril persistence and width on nanopatterns compared to flat surfaces, indicating enhanced higher-order organization. Thus, myofibril maturation in the D65A mutant on nanopatterns is more similar to WT, demonstrating that external cues (provided by nanopatterns) can overcome internal cues (loss of contraction) and partially rescue structural defects in non-contractile cardiomyocytes.

**Figure 5.**
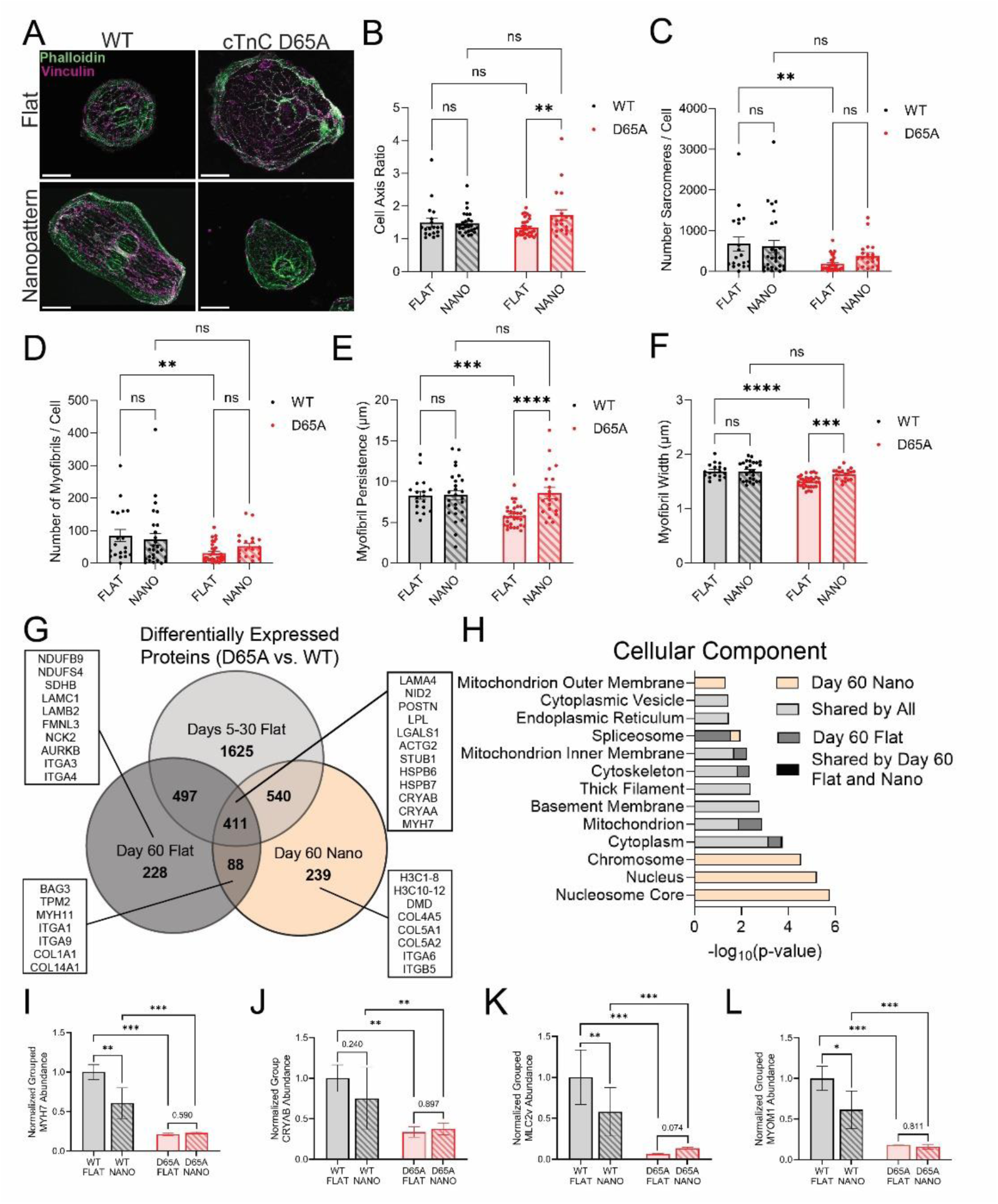
Nanopatterns promote structural maturation in cTnC D65A cardiomyocytes. WT and cTnC D65A cardiomyocytes were replated onto flat or nanopatterned surfaces on Day 46 and analyzed by immunofluorescence microscopy on Day 60 **(A-F)**. **A)** Representative images of wildtype and mutant cardiomyocytes labeled with phalloidin (green) and vinculin antibody (purple). Scale bars = 30 µm. **B-F)** Quantification of cell axis ratio **(B)**, sarcomere number per cell **(C)**, myofibril number per cell **(F)**, myofibril persistence (the length over which myofibrils remain straight) **(E)**, and myofibril width **(D).** Significance was determined via 2-way ANOVA with multiple comparisons. **G-L)** WT and cTnC D65A samples from flat and nanopatterned surfaces on Day 60 were analyzed by mass spectrometry. **(G)** Venn diagram showing the number of shared and unique proteins that were differentially expressed between WT and mutant for different times and conditions. Examples from each group are shown in boxes. **H)** Enriched Cellular Component GO terms for subsets of differentially expressed proteins. Groups were defined based on Panel G. **I-L)** Grouped abundances of maturation markers MYH7 **(I)**, CRYAB **(J)**, MLC2v **(K)**, and MYOM1 **(L)**. Values were normalized to WT plated on flat surfaces. Statistics were performed by Proteome Discoverer (Table S1 and Table S3). Error bars show the standard error of the mean.

To investigate how contractility affects ECM sensing, we quantified focal adhesion (FA) geometry on flat surfaces and nanopatterns by staining for vinculin. FAs are proteinaceous complexes that physically link the cytoskeleton to the ECM, transmit mechanical tension, and act as signaling hubs for ECM sensing.^35^ We observed no change in average FA size between any group (**Figure S6A**). However, the D65A cells had more elongated FAs on nanopatterns compared to flat surfaces, whereas WT did not undergo this shift (**Figure S6B**). In addition, while both WT and D65A cardiomyocytes had more aligned FAs on nanopatterns than flat surfaces, D65A FAs on nanopatterns were more aligned than their WT counterparts (**Figure S6C**). These differences suggest that loss of contractility changes FA geometry and may facilitate ECM sensing.

To identify other nanopattern-specific changes between mutant and WT cardiomyocytes, we grouped proteins that were differentially expressed in D65A vs. WT on the nanopatterns, excluding proteins that were also differentially expressed on flat surfaces at any timepoint (**Figure 5G**). This group was highly enriched in proteins that localize to the nucleus and subnuclear structures like the nucleosome core and chromosomes (**Figure 5H**), suggesting that internal mechanics like sarcomere contraction influence ECM-nucleus crosstalk. However, when we stained nuclei with Hoechst 33342, we found no difference in nuclear area, nuclear elongation, or nuclei number per cell between WT and D65A cardiomyocytes (**Figure S7A-C**). Together, the data suggests that contractility alters the nuclear proteome, but these changes do not affect gross nuclear morphology. In contrast, surface type did modify nuclear morphology, with nanopatterns inducing smaller nuclei in WT and D65A cells. This implies that both contractile and non-contractile cardiomyocytes sense nanopatterns and transmit these external mechanical cues to the nucleus.

### Nanopatterns Induce Differential Maturation Responses in Wildtype and cTnC D65A Cardiomyocytes

We next probed how nanopatterns affect maturation of the proteome. D65A cells on nanopatterns had reduced maturation marker abundance compared to WT (**Figure 5I-L**), recapitulating the phenotype observed on flat surfaces. There was no significant difference between D65A cardiomyocytes on flat and nanopatterned surfaces, but WT cells on nanopatterns surprisingly had a significant decrease in 3 out of the 4 markers (MYH7, MLC2v, and MYOM) compared to WT on flat surfaces (**Table S3**). This suggests that nanopatterns disrupt maturation of the WT proteome.

To investigate this idea further, we next analyzed how nanopatterns influence the expression of ≥60 OXPHOS proteins (**Figure 6A**). In WT cardiomyocytes, 15% of the proteins were significantly decreased on nanopatterns, whereas only 5% were significantly increased. However, in the mutant, ∼25% of our OXPHOS proteins were increased on nanopatterns compared to ∼2% that were decreased. The GO-terms “respiratory chain” and “electron transport” were also enriched among proteins upregulated by nanopatterns in D65A cardiomyocytes (**Figure 6B**). These findings demonstrate that nanopatterns increase expression of OXPHOS components in non-contractile cardiomyocytes but decrease them in contractile, WT cardiomyocytes.

**Figure 6.**
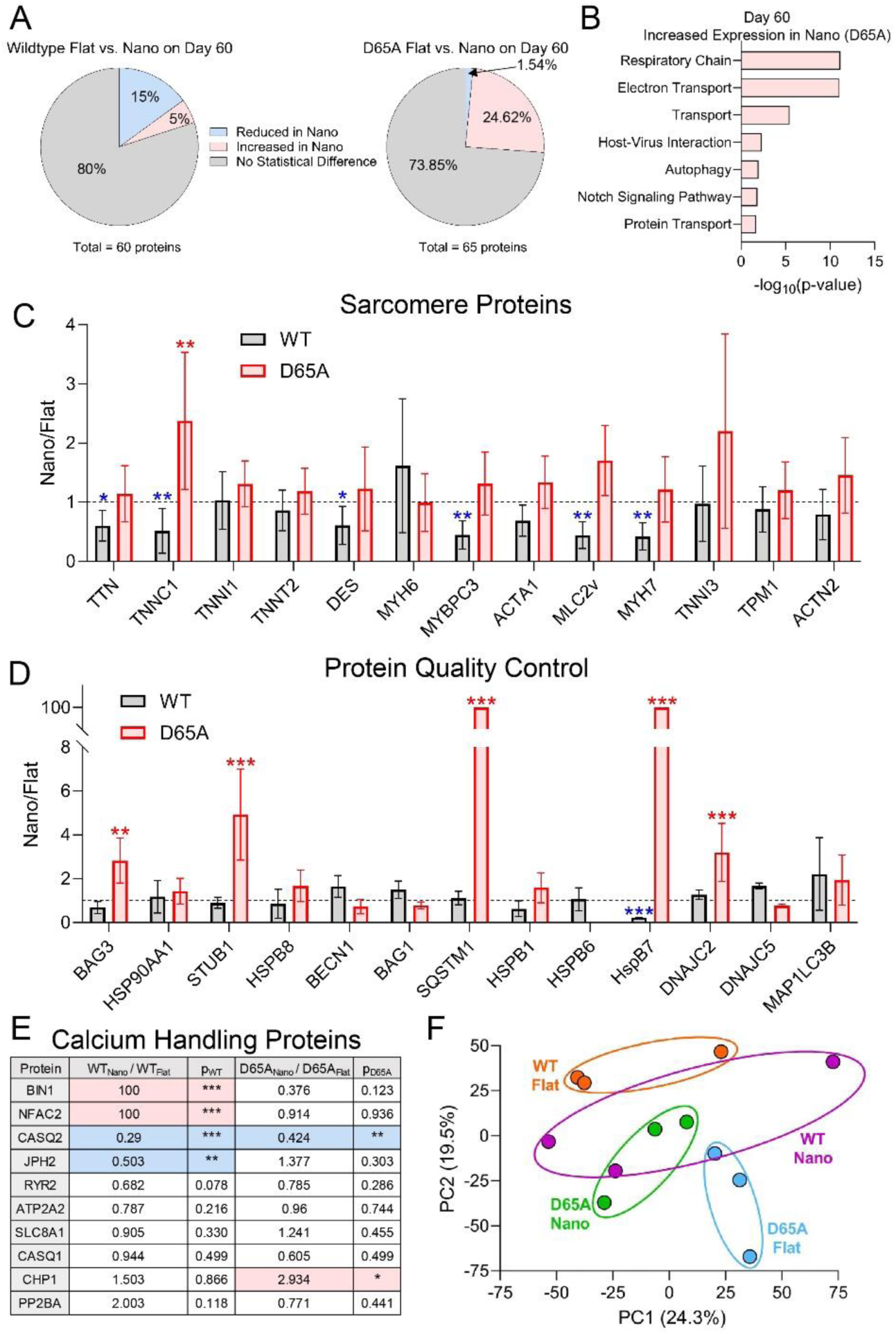
Nanopatterns induce differential responses in wildtype and cTnC D65A cardiomyocytes. **A)** Pie chart showing percentages of oxidative phosphorylation proteins from our panel that were significantly different on nanopatterns versus flat surfaces for WT (left) and cTnC D65A (right) cardiomyocytes on Day 60. **B)** Biological Process GO terms enriched among proteins that were upregulated on nanopatterns in D65A cardiomyocytes. **C-E)** Abundance ratios (Nano/Flat) for sarcomeric **(C)**, protein quality control **(D),** or calcium handling **(E)** proteins for WT and cTnC D65A cardiomyocytes. Dotted line marks a ratio of 1. Significant differences between nanopatterns and flat surface groups are indicated by asterisks (*<0.05, **<0.01, ***<0.001). Color indicates increased (red) or decreased (blue) expression on nanopatterns. **F)** Principal component analysis of WT and cTnC D65A samples on nanopatterns and flat surfaces on Day 60. Individual samples are represented by a dot. Colors and ovals indicate samples from the same group.

Nanopatterns also had differential effects in D65A versus WT cardiomyocytes in other maturation-related contexts. Several sarcomere proteins (including TTN, TNNC1 (cTnC), MYBPC3, and DES) were reduced in WT cardiomyocytes on nanopatterns compared to their flat surface counterparts (**Figure 6C**). In contrast, nanopatterns did not generally modulate sarcomere protein levels in D65A cells; only cTnC (TNNC1) was upregulated on the nanopatterns. PQC protein levels were largely unchanged in WT but upregulated in D65A cardiomyocytes on nanopatterns (**Figure 6D**). Calcium handling proteins showed no clear trend in nanopattern-induced changes in either mutant or WT (**Figure 6E**). Overall, nanopatterns had a negative or neutral effect on WT structural and proteomic maturation yet a neutral or positive effect on D65A maturation.

Principal component analysis encapsulates some of the trends we observed in the nanopattern data (**Figure 6F, Table S4**). D65A cells on nanopatterns cluster more closely with WT compared to D65A cells on flat surfaces. However, nanopatterns also cause WT to shift toward the immature and non-contractile mutant samples. Taken together, these data suggest that nanopatterns facilitate structural and metabolic maturation in non-contractile cardiomyocytes, but in WT cells, nanopatterns reduce proteomic maturation markers. These differences imply that external mechanical cues like nanopatterns are only beneficial in immature or contractility-deficient cells.

## Discussion

To investigate the role of internal and external mechanical cues on iPSC-derived cardiomyocyte development, we leveraged the cTnC D65A mutation to block calcium-activated contractility (an internal mechanical signal) and employed nanopatterned surfaces (an external mechanical signal). Surprisingly, we found that sarcomeres still form in the D65A cells, though these sarcomeres are underdeveloped and disorganized. Mutant cells also exhibit altered calcium transients (increased calcium release, abnormal kinetics) and significant maturation defects at the proteomic level. Plating D65A cardiomyocytes on nanopatterns improved maturation metrics, leading to more elongated cell morphologies, enhanced higher-order myofibril structure, and upregulated OXPHOS machinery. On the other hand, nanopatterns did not induce structural changes in the WT cardiomyocytes and led to the downregulation of some sarcomeric and metabolic maturation markers.

Previous work by others concluded that myosin-mediated force generation^7^, TnT^36^, and TnI^37^ are required for sarcomere formation in striated muscle. These studies led us to initially predict that the cTnC D65A mutant would prohibit sarcomere assembly. Instead, the ability to produce sarcomeres was clearly preserved. This finding highlights our incomplete understanding of the exact mechanisms of sarcomerogenesis. The prevailing model (known as the pre-myofibril model) suggests that sarcomere formation begins with the assembly of muscle-specific stress fibers at the cell periphery.^6–9^ These stress fibers migrate into the cell interior, shifting their protein composition and developing into myofibrils. Troponin and sarcomeric tropomyosin are present in pre-myofibrils,^38^ and they are thought to be incorporated into filaments immediately after actin polymerizes.^39^ It is not fully understood how muscle-specific stress fibers evolve into myofibrils, but some evidence suggests that they may act like a template. Our data indicates that the absence of contraction does not preclude the transition of pre-myofibrils into myofibrils but that further myofibril maturation is impaired.

Sarcomere formation in iPSC-derived cardiomyocytes is blocked by treatment with the myosin-II ATPase inhibitor blebbistatin, suggesting that myosin binding to thin filaments and generating force is somehow necessary for sarcomerogenesis.^7^ Taken with our data, the evidence points to the need for some degree of unregulated actin-myosin interaction (i.e., not dependent on troponin/tropomyosin). One possibility is that these interactions come from non-muscle myosins, as the gatekeeping function of tropomyosin, and therefore troponin, is dependent on myosin isoform.^40^ Supporting this idea, evidence from *in vitro* motility assays suggests that non-muscle myosins (NMMIIA, NMMIIB, NMMIIC) are not inhibited by striated muscle α-tropomyosin, thus circumventing the need for a functional cTnC.^41^ Additionally, pan-inhibition of myo-II isoforms with blebbistatin abolishes sarcomere assembly,^7^ but specific skeletal muscle myo-II inhibition allows sarcomeres to form.^10^ The involvement of non-muscle myosins in sarcomere assembly is debated,^7^ but some data suggests that non-muscle myosins localize to pre-myofibrils^42^ and play a role in sarcomerogenesis.^6^

Unlike TnC Ca^2+^ binding, the other two components of the troponin complex (TnT and TnI) are required for sarcomere assembly.^36,37^ The different outcomes may reflect the distinct functional roles of each subunit or perhaps a requirement for the physical troponin proteins but not their full functionality. Troponin I inhibits myosin binding, and its loss in drosophila indirect flight muscles prevents sarcomere formation, perhaps from uncontrolled myosin activity.^37^ Troponin T stabilizes tropomyosin onto the thin filament,^36^ and depletion of all three isoforms in zebrafish skeletal muscle leads to tropomyosin displacement, again disrupting sarcomere formation via uncontrolled myosin activity. Notably, sarcomerogenesis is compromised by both uncontrolled myosin activity and myosin inhibition but blocking only calcium-activated (troponin-dependent) myosin activity is tolerated. As such, we speculate that specific inhibition of TnI’s switch function would still permit sarcomere assembly.

Although not required for sarcomerogenesis, sarcomere contraction clearly facilitated maturation. The D65A mutant had fewer, smaller, and more disorganized myofibrils than WT. Corroborating the structural data, D65A’s proteomic markers also suggested defective maturation. Our data aligns well with findings in *Xenopus tropicalis* embryos demonstrating the need for skeletal muscle myo-II activity for sarcomere maturation.^10^ This previous study also found that tension is required for organizing both thick filaments and z-discs. These effects may be the basis for the maturation abnormalities we observed in cTnC D65A cardiomyocytes.

Interestingly, although proteomic maturation markers were reduced in D65A cardiomyocytes compared WT on Days 30 and 60, they were often upregulated at earlier timepoints. These early samples may include some cells that failed to differentiate because they were collected prior to cardiomyocyte purification (Day 20). However, given that cTnC is not expressed in undifferentiated cells and that there is no ostensible difference in differentiation efficiency between the mutant and the WT (**Figure S3**), it is likely that the changes in maturation markers come from the cardiomyocyte population. These elevated protein levels may be a response to the loss of contractility and an attempt to develop structures that facilitate contraction, representing an example of cardiomyocyte mechanosensing of internally generated forces.

A key finding of our study was that nanopatterned surfaces enhance aspects of maturation (myofibril structure, morphology, OXPHOS) in D65A cardiomyocytes, partially compensating for the absence of the internal cue of sarcomere contractility. This contrasts with our findings in contractile cardiomyocytes, where nanopatterns did not significantly alter structural metrics and reduced expression of maturation-associated sarcomeric and OXPHOS proteins. This outcome was unexpected because previous studies suggest that nanopatterns are beneficial for WT maturation. Nanotopography was first hypothesized to improve the characteristics of iPSC-CMs by Kim et al (2009),^43^ who observed rat cardiomyocytes aligning with the underlying grooves in the extracellular matrix in SEM micrographs. They constructed nanopatterned surfaces and found that neonatal rat ventricular myocytes aligned along its grooves. The nanopatterns also led to increased expression of the cell-cell coupling protein connexin 43, faster action potential conduction velocities, and a more uniform contraction direction. Other groups later showed that plating iPSC-derived cardiomyocytes on nanopatterns increases cell area^44,45^, perimeter^45^, and sarcomere length^45^, while reducing circularity,^45,46^ all of which are indicative of enhanced maturation. Nanopatterns also increase mRNA levels of maturation markers (e.g., MYH7, RYR2, TNNI3 (cTnI))^46,47^ and enhance maximum contraction velocity,^47^ though slower contraction velocities on nanopatterns have also been detected.^44^

A major advantage of the current study is that we obtained a broad overview of the proteome rather than analyzing mRNA transcript levels for a limited number of maturation markers. Importantly, mRNA levels do not equate to protein levels due to regulatory controls involving mRNA translation and protein degradation.^48^ Thus, it is important to specifically evaluate the proteomic effects of nanopatterns on iPSC-CM maturation, which, to our knowledge, had not been previously assessed.

The downregulation of sarcomeric and OXPHOS proteins in WT cardiomyocytes suggest that nanopatterns hindered WT maturation. There are key differences between our study and previous work that may explain why nanopatterns had this effect. Other groups cultured cardiomyocytes for shorter time periods on flat surfaces before replating onto nanopatterns and also analyzed their samples at overall earlier timepoints (**Table S5**).^43,45–47^ In contrast, we cultured our cells for 46 days prior to replating onto nanopatterns and continued to mature the cardiomyocytes until Day 60. The prolonged culture time—both in total and prior to replating onto nanopatterns—likely enabled our cardiomyocytes to reach a more advanced maturation state. Our results may mean that nanopatterns are not beneficial if cardiomyocytes are introduced to them after a certain stage of maturation, with only immature cardiomyocytes being susceptible to/tolerant of nanopattern-mediated changes. Alternatively, nanopatterns may only exert negative effects at advanced stages of maturation while facilitating maturation at earlier stages. Given that nanopatterns enhanced several maturation metrics in immature D65A cells but not in mature WT cells, our results support both explanations. Yet, the differential response to nanopatterns between WT and D65A cardiomyocytes could also be rooted in contraction itself rather than contractility’s downstream effect of altered maturity. Since we replated our cells onto nanopatterns after differences in maturity had already manifested, we cannot distinguish between these possibilities.

Although disrupted WT maturation on nanopatterns is the simplest explanation for our results, other explanations exist and should be considered. It is possible that decreased sarcomeric and OXPHOS protein levels do not indicate reduced maturity but instead reflect differences in protein abundance regulation. For example, some groups have suggested that sarcomeric proteins are held in cytoplasmic pools prior to incorporation into the sarcomere and after removal.^49,50^ If nanopatterns reduce the size of these pools in WT cardiomyocytes— perhaps through the stabilization of myofibril structure and subsequent decrease in protein turnover—then this would lead to a reduction in sarcomere protein content and potentially lower the demand for OXPHOS proteins, as we observed. However, the existence of this cytoplasmic pool is debated, with recent evidence indicating that only newly translated proteins are added to the sarcomere while removed proteins are immediately degraded.^51^ Regardless of the underlying mechanism, our results highlight the need for further investigation into the effects of nanopatterns on maturation and sarcomeric protein abundance.

Our findings also have implications for cardiac regeneration. Recently, transplantation of iPSC-derived non-contractile cardiomyocytes in rats after ischemia/reperfusion injury was shown to preserve function similarly to contractile cardiomyocytes.^52^ The improvement of D65A cardiomyocytes on nanopatterns in our study suggests that ECM cues within the myocardium could facilitate maturation of transplanted non-contractile cardiomyocytes *in vivo*.

This study provides novel insight into cardiomyocyte development by establishing the dispensability of calcium-activated contraction in sarcomerogenesis and its importance in cardiomyocyte maturation. We also demonstrated that nanopatterns improve maturation in immature, non-contractile cardiomyocytes, suggesting that external mechanical cues may partially compensate for defective contractility, like that which occurs often in cardiovascular disease. However, further study is required to elucidate the mechanisms behind this integration of internal and external cues. Understanding such biological programming will be crucial for optimizing the utility of iPSC-cardiomyocytes and ultimately moving them from the lab into the clinic.

## Acknowledgements

Schematic of experimental workflow was created using BioRender (Delligatti, C. (2024) https://BioRender.com/l04l280).

